# Sickness behaviour reduces network centrality in wild vampire bats

**DOI:** 10.1101/2020.03.30.015545

**Authors:** Simon P. Ripperger, Sebastian Stockmaier, Gerald G. Carter

## Abstract

Sickness behaviours, like lethargy, can slow the spread of pathogens across a social network. We conducted a field experiment to investigate how sickness behaviour reduces individual connectedness in a high-resolution dynamic social network. We captured adult female vampire bats (*Desmodus rotundus*) from a wild roost. To create ‘sick’ bats, we injected a random half of the bats (n=16) with the immune-challenging substance, lipopolysaccharide, and injected control bats with saline (n=15). Over the next three days, we used proximity sensors to continuously track their associations under natural conditions. The ‘sick’ bats showed a clear decrease in social connectedness (degree, strength, and eigenvector centrality). Bats in the control group encountered fewer ‘sick’ bats and also spent less time near them. These effects varied by time of day and declined over 48 hours. High-resolution proximity data allow researchers to define network connections based on how a pathogen spreads (e.g. the minimum contact time or distance for transmission). We therefore show how the estimate of the sickness effect changes as network ties are defined using varying distances and durations of association. Tracking the effects of sickness behaviour on high-resolution dynamic social networks can help create more sophisticated simulations of pathogen transmission through structured populations.

## Background

As a pathogen spreads across a population, sickness behaviours – like lethargy, increased sleep, and reduced movement – can slow pathogen spread, because less socially connected individuals are often less likely to transmit a pathogen [1-3]. This sickness-induced ‘social distancing’ can be important for modelling pathogen transmission as a social network changes over time (i.e. a dynamic social network [4]). Tracking the effects of sickness behaviour on a dynamic social network requires large datasets with temporal and spatial resolutions that are high enough to be ecologically useful. Automated tracking of animal associations typically occurs in the lab [1] or at specific field locations such as feeders or nest boxes [2]. Proximity sensors by contrast can measure association times and durations, at high spatial and temporal resolution, among free-ranging animals at any location [5]. Proximity tracking is therefore a potentially powerful tool for understanding how individual sickness behaviour reshapes a social network.

Here, we induced sickness behaviour in wild-caught vampire bats using injections of lipopolysaccharide (LPS), which mimics the symptoms of a bacterial infection without an active pathogen. LPS treatments allow us to isolate the effects of sickness behaviour from parasite-specific manipulations of host behaviour [6, 7]. After injections, we tagged both the ‘sick’ bats (injected with LPS) and control bats (injected with only saline) with proximity sensors [8]. We released them back into their wild colony and tracked changes in their association rates. Based on the effects of LPS on the physiology and behaviour of captive vampire bats [6, 7], we predicted reduced association rates between ‘sick’ bats and control bats in the wild.

Indeed, LPS-induced sickness behaviour caused a dramatic decrease in network centrality. The control bats encountered fewer ‘sick’ individuals and also spent less time near them. For studying pathogen transmission, the links (or edges) in a social network would ideally be defined based on the pathogen-specific transmission mechanism, because some pathogens require longer or closer physical contact. In practice, however, most social network edges are defined based on technological or statistical limitations. Therefore, we used resampling to show how estimates of the sickness effect change when network ties are defined using varying distances or durations of association.

## Methods

### Inducing sickness behaviour

We captured bats from a colony of common vampire bats (*Desmodus rotundus*) inside a hollow tree at Lamanai, Belize. Before sunset on April 24^th^ 2018, we blocked all exits of the roost except one and we used a handnet and mist nets to capture about 100 vampire bats (including 41 females) until 0500 h the next morning. We kept females in cotton cloth bags, and measured their mass to ensure they did not differ between the randomly assigned treatment and control injections (difference in mass = 0.17 g [95% CI: −2.9, 2.4]). We randomly assigned the females to the test or control treatment by flipping a coin, then adjusted to ensure more balanced samples. We injected the individuals in the test group under the dorsal skin with 70-100 µl of LPS (lipopolysaccharide in phosphate-buffered saline, L2630 Sigma-Aldrich, St Louis, MO, U.S.A.) at a dosage of 5 mg/kg, following previous studies with this species [6, 7]. Bats in the control group received an injection of the same volume per body mass of phosphate-buffered saline. One hour after injection, we released 34 females, tagged with proximity sensors, back into their roost.

### Proximity tracking

To track dyadic associations among the bats, we used custom-built proximity sensors (see [5, 8, 9] for details). The sensors weighed 1.8 g (including battery and housing) and were glued to the dorsal fur using skin-bonding latex adhesive (Montreal Ostomy Skin-Bond). Tag weights were 4.5 – 6.9 % of each bat’s mass, in accordance with recommendations for short-term tracking of bats [10]. We placed the antennas of a base station inside the roost for encounter data download. Each encounter observation includes a duration and received signal strength indicator (RSSI), which can be used as an estimate for a minimum distance between two tagged bats during the encounter. We defined a ‘proximity index’ as the percent quantile of all RSSI values. To define association, we used a proximity index of 85% (i.e. the top 15% of all encounters ranked by signal strength, Fig. S1). We chose this value by using the same RSSI value as a previous study linking wild associations to captive interactions (−27 dbm [8]). Past work [8] suggests these associations involve a proximity of about 0-50 cm.

We excluded data from three sensors, which apparently dropped off the bat, either inside (n=2) or outside (n=1) the roost, evident from the sensor’s constant contact with the base station (i.e. no evidence of exiting or entering the roost). We therefore used association data from 16 ‘sick’ bats and 15 control bats.

### Network construction

We created social networks where edges were association time. To track associations over the day, we created social networks for each hour. To measure an LPS effect size, we created a network for the entire period where we expected an LPS effect based on past work [7]. This “treatment period” was 3 to 9 h post-injection (1700 - 2300 h). We did not include associations from the second half of the night because we observed, visually and in the sensor data, that most of the bats left the roost to forage after midnight. For comparison, we also created two more networks for the corresponding times of day (24 and 48 h later).

### Hypothesis-testing

To test the effect of LPS on three measures of network centrality, we first fit a general linear mixed-effects model with treatment (LPS, saline) and day (1, 2, 3) as fixed effects, bat as a random effect, and the network centrality measure (degree, strength, and eigenvalue, respectively) as a response. We then extracted the standardized model coefficients for the treatment effect and the interaction between treatment effect and day. If we detected an interaction, we also fit a linear model for the observations within the first and last day separately and extracted those standardized treatment effect coefficients. To get two-sided p-values, we created 10,000 null datasets where the treatment was re-assigned randomly among bats at the start of the study, then measured the proportion of the null coefficients that were greater than the observed coefficients, and then doubled those one-sided p-values. This procedure creates a null model accounting for the non-independent and non-normal structure of the network data [11]. To assess assortativity of sick and control bats over time, we calculated for each hour the association probability (proportion of possible pairs that were associated) and the mean association time (total seconds per period) for three dyad types: control-control, control-sick, and sick-sick. For all 95% confidence intervals, we used basic nonparametric bootstrapping with 5,000 iterations.

### Measuring effects of network construction on effect size of sickness behaviour

Defining network edges requires deciding what minimum proximity or duration constitutes an ‘association’. With proximity sensor data, this definition is flexible but requires a trade-off between maximizing the sample size of observations and filtering for observations that are more meaningful (e.g. closer proximities or longer durations). We inspected how the size and precision of the LPS effect changed with variations in how networks were constructed. To do this, we resampled our data using different definitions of association, then we plotted changes in the number of observations and the treatment effect size (defined as the unstandardized model coefficient during the treatment period). As a measure of relative detectability, we used the p-value from the parametric linear model. To investigate the effect of minimum encounter duration, we defined association at one proximity index (85% as in our original analysis), but filtered encounters using several values of minimum duration that varied from 0 - 1200 s. To investigate the effect of minimum encounter proximity, we set no minimum duration (as in our original analysis), but filtered encounters using a proximity index threshold that varied across several values from 76 - 98 %. Proximity index thresholds below 76% are not informative given the roost size; thresholds over 98% use less than 2% of the data. Finally, to assess if the LPS treatment effect size was robust across different proximity thresholds, when controlling for number of observations, we used several proximity index thresholds varying from 75% to 94% but we randomly sub-sampled the same number of observations in each case (5,258 encounters or 95% of the number of observations at the 94% proximity index threshold). For each proximity index threshold, we obtained 200 effect size estimates with different random sub-samples.

## Results

### LPS-induced sickness behaviour reduces association rates

Compared to the control group, ‘sick’ (LPS-injected) bats associated with fewer groupmates (lower degree centrality), spent less time with them (lower strength centrality), and were less socially connected to the entire network (lower eigenvector centrality), and these effects diminished with time (Figure 1, Table 1). During the hours of the treatment period, a control bat had on average a 49% chance of associating with each control bat ([95% CI: 44, 54], n = 105 pairs), but only a 35% chance of associating with each ‘sick’ bat ([31, 38], n = 240 pairs). During the treatment period, the mean association rate for two control bats was 15 min per h [13, 18], but for a control and ‘sick’ bat, it was only 10 min per h ([8, 11], Figure 2).

**Table 1.**
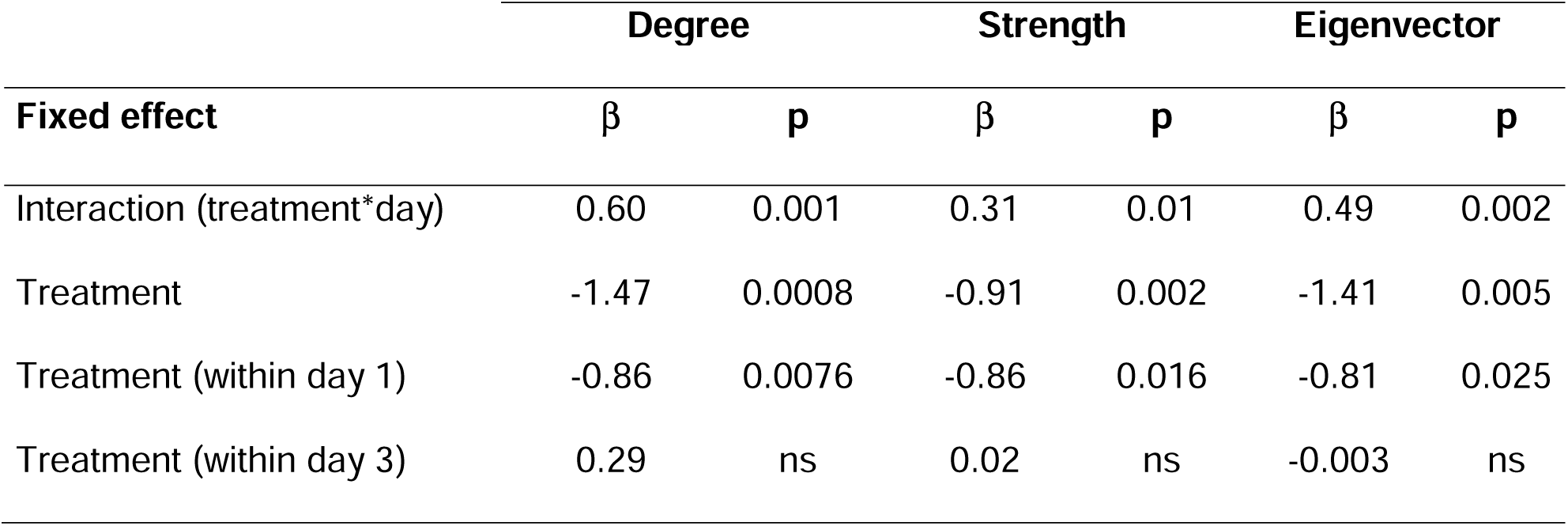
Model estimates of LPS treatment on network centrality. Standardized coefficients (β) and two-sided p-values are reported for degree, strength, or eigenvector centrality.

|  | Degree |  | Strength |  | Eigenvector |  |
| --- | --- | --- | --- | --- | --- | --- |
| Fixed effect | $\beta$ | p | $\beta$ | p | $\beta$ | p |
| Interaction (treatment*day) | 0.60 | 0.001 | 0.31 | 0.01 | 0.49 | 0.002 |
| Treatment | -1.47 | 0.0008 | -0.91 | 0.002 | -1.41 | 0.005 |
| Treatment (within day 1) | -0.86 | 0.0076 | -0.86 | 0.016 | -0.81 | 0.025 |
| Treatment (within day 3) | 0.29 | ns | 0.02 | ns | -0.003 | ns |

**Figure 1.**
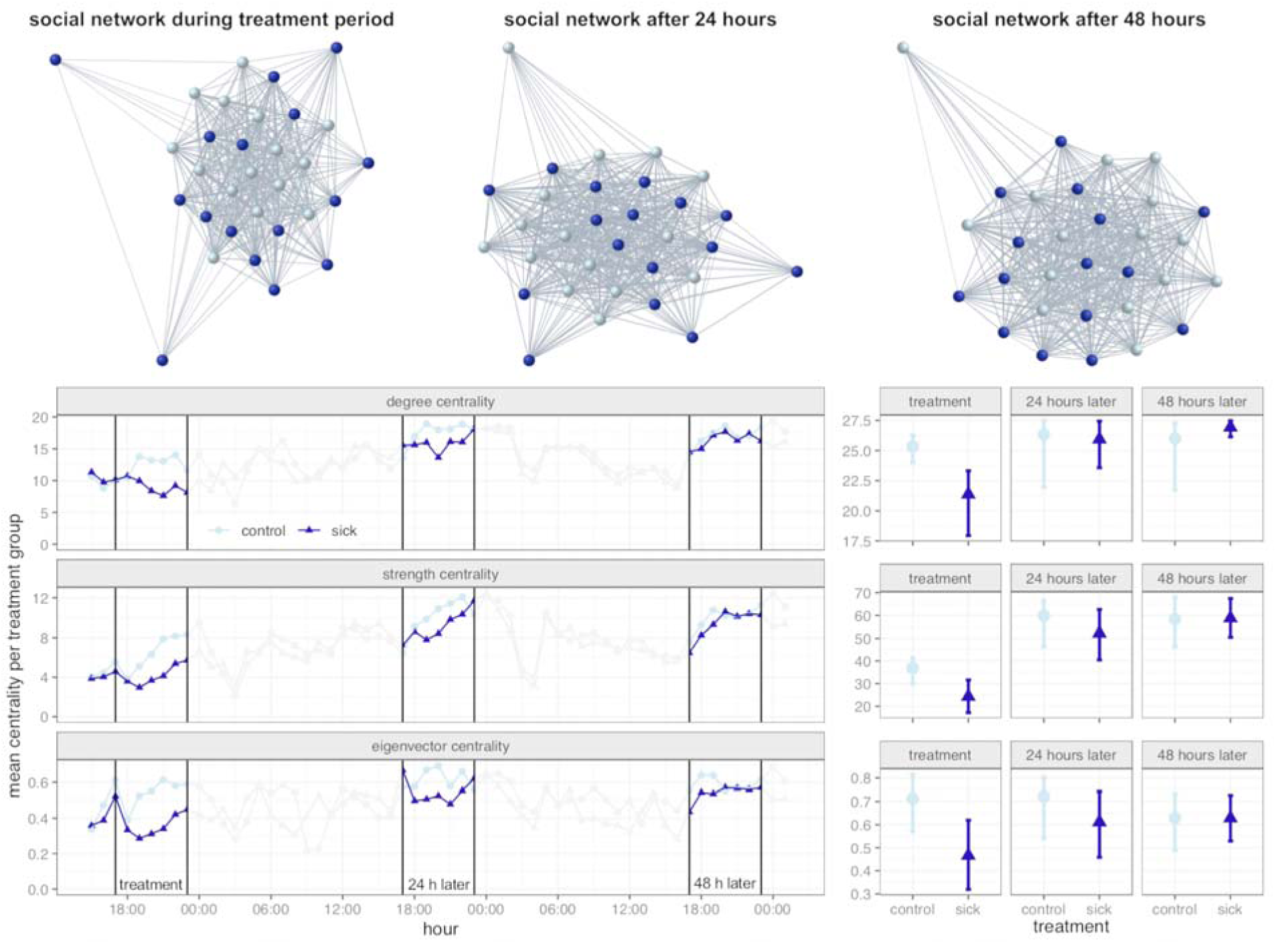
Centrality decreased in LPS-injected bats. ‘Sick’ LPS-injected bats (dark nodes) were less socially connected than control bats (light nodes) and this effect diminished over 48 hours. Edge weights are association times (log_10_-transformed). Spatial positions are based on the graph embedder (GEM) force-directed layout algorithm. Left-hand time-series panel shows that three measures of mean centrality (degree, strength, and eigenvector) were lower in the ‘sick’ test group (dark triangles) compared to the control group (light circles). Solid vertical lines show the treatment period and the corresponding hours on the next two days. Right-hand panels show the centrality measures of each group during the entire treatment and post-treatment periods. Centrality values in the right-hand panel are higher because networks were constructed for the whole period. Values for strength centrality are hours rather than seconds. Error bars are bootstrapped 95% confidence intervals.

**Figure 2.**
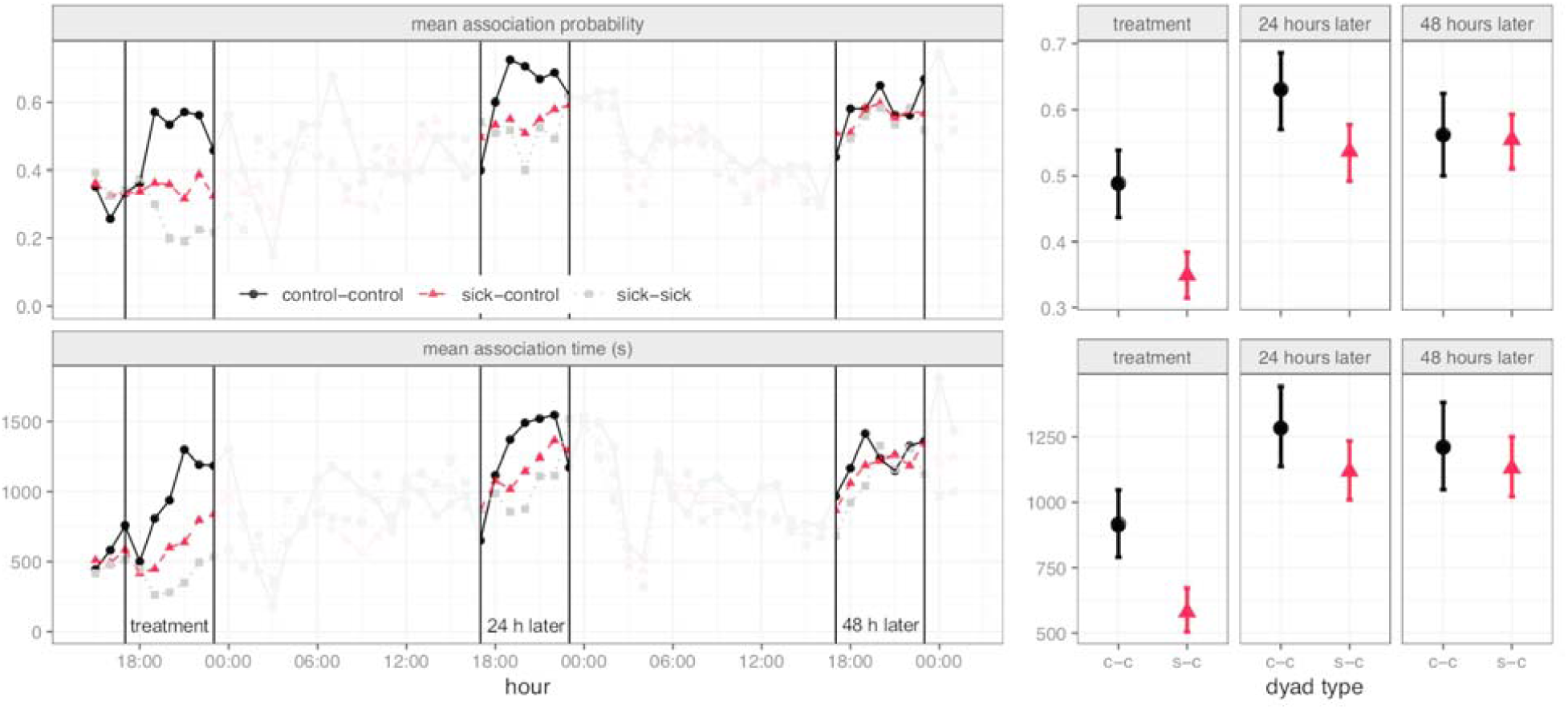
Saline-injected control bats associated less with LPS-injected ‘sick’ bats. The hourly association probability (top) and mean association time (bottom) are shown for two control bats (black circles), two sick bats (grey squares), and one of each (red triangle). Panel B shows the hourly probability (top) and mean time (bottom) of association between a control bat and another control bat (black circles) or a control and a ‘sick’ bat (red triangles), during each period. Error bars are bootstrapped 95% confidence intervals.

### Effect sizes and detectability of sickness effects depend on network construction

When we increased the minimum threshold of time defining an association, we observed that the estimate of the treatment effect grew larger, but eventually became smaller and less clear as the number of observations decreased (Figure 3A). When only using associations over 15 min, the effect became harder to detect. Next, when we increased the minimum proximity index threshold defining an association, we again observed that the treatment effect estimate grew larger but less clear as the number of observations declined (Figure 3B). When controlling for the number of observations, the treatment effect was relatively stable across different proximity thresholds (Figure S2).

**Figure 3.**
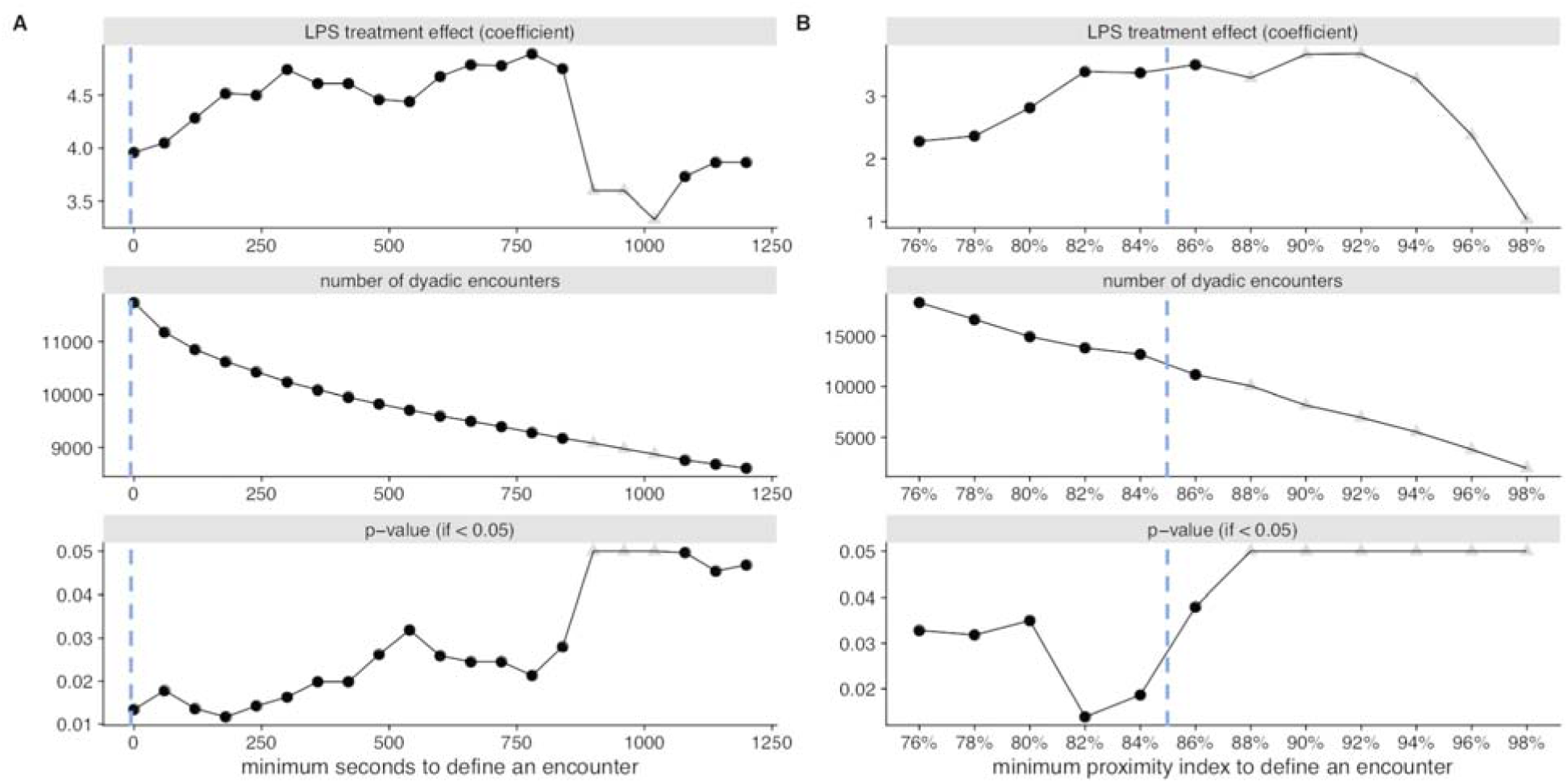
Network construction alters the magnitude and detectability of treatment effects on degree centrality. Association rates are based on many encounters that can be defined by a minimum duration (A) or proximity (B). Panel A shows how the minimum threshold of time to define a social encounter (x-axis) affects the estimate of the treatment coefficient (top), the sample size of dyadic encounters (middle), and the p-value for the parametric model (cropped at 0.05). Panel B shows how the same measures are affected when social encounters are defined using a different minimum estimate of relative spatial proximity. Dashed lines shows duration and proximity index thresholds used in main analysis. Grey triangles indicate estimates with parametric p-values >0.05. Note that this p-value is based on violated assumptions of normality and independence, so it should be used only as a proxy for relative detectability, not for actual inference.

## Discussion

Sickness effects, induced by LPS injections, decreased the network connectivity of ‘sick’ bats (Figure 1), and reduced the probability of encounters and the association times between female vampire bats in the ‘sick’ and control group (Figure 2). These effects were not caused by spatial assortativity of ‘sick’ bats, because associations between two ‘sick’ bats were even lower (Figure 2). The behavioural changes causing these effects are evident from captive studies showing that LPS-injected vampire bats are less active [7]. They also produce fewer contact calls [12], which attract bonded partners [13]. When tested in a flight cage, ‘sick’ vampire bats engaged in social grooming with fewer partners [6], but we observed dramatic reductions in social grooming even when captive pairs were forced into close association [7]. LPS-induced sickness behaviours therefore reduce both associations and behaviours, like social grooming, which can further enhance pathogen transmission between associated bats.

The effects of LPS vary by dose and among species [14]. For instance, in LPS-injected rats, social exploration of juvenile conspecifics and locomotor activity largely returned to normal after 24 hours [15]. Here, we found evidence for behavioural effects after 24 hours, which could be due to an ongoing immune challenge, exhaustion post-recovery, or an attempt to save energy from not foraging on the previous night. The observed pattern could also result from control bats avoiding the test group based on past interactions, but captive studies on LPS effects have not observed clear evidence for avoidance behaviour [6, 7].

Restructuring of social networks following an infection can occur through four nonmutually exclusive processes. First, as we observed here, infection-induced lethargy can passively reduce associations (mice: [2], humans: [3]). Second, individuals might actively avoid contact with infected conspecifics (e.g. lobsters: [16], bullfrog tadpoles: [17], mice: [18], mandrills: [19]). Third, individuals might actively and collectively restructure their social structure (eusocial insects: [1, 20]; humans: [21]). Fourth, parasites can manipulate host behaviour to restructure host networks in favour of parasite transmission [22, 23]. These processes can also interact. For example, infection-induced changes in host behaviour can induce feedbacks that alter parasite manipulation behaviour on both developmental and evolutionary timescales [22].

Depending on the goals of a study, sickness behaviour or pathogen transmission can be measured or modelled at varying spatial and temporal scales. Studies using passive integrated transponder tags to track free-ranging mice showed a decreased probability of sharing a nestbox [2]. Here, proximity sensors allowed us to continuously measure proximity, even within a single roost. On a larger spatial and temporal scale, pathogen transmission crucially depends on movements between roosts and sites (e.g. rabies in vampire bats [24]). Conceptually or mathematically, the social network of transmission rates between individuals within each site can be embedded within a single node of a larger network mapping transmission rates between sites (e.g. [25]).

When defining network edges with continuous proximity data, there is a trade-off between the number of observations and filtering closer encounters that are more relevant for a given behaviour or pathogen. In this study, we used resampling to show the effect of defining association at various thresholds of minimum duration and distance (Figure 3). We recommend this resampling procedure for testing the robustness of an effect across the range of durations or distances that are biologically meaningful (Figure S2). As tracking technology improves the capacity to create dynamic networks from massive, high-resolution datasets, we expect researchers to gain transformative insights into the patterns and processes underlying the spread of pathogens, information, or behavioural states.

## Ethics statement

This work was approved by the Institutional Animal Care and Use Committee and the American Museum of Natural History (Protocol # AMNHIACUC-20180123).

## Data and code availability

The datasets and R code for this article can be found at Figshare: https://doi.org/10.6084/m9.figshare.12045450.v2

## Acknowledgements

We thank Michelle Nowak for help with fieldwork and data collection. We thank Brock Fenton, Nancy Simmons, and Dan Becker for help with logistics and permits. We thank Frieder Mayer for funding and Björn Cassens and Niklas Duda for technological support.

## Author contributions

GGC conceived of the study. SPR and GGC participated in fieldwork, data collection, analysis, and writing. SS created the LPS treatments and participated in writing. All authors gave final approval for publication and agree to be held accountable for the work performed therein.

## References

1. Stroeymeyt N., Grasse A.V., Crespi A., Mersch D.P., Cremer S., Keller L. 2018 Social network plasticity decreases disease transmission in a eusocial insect. Science 362(6417), 941–945.

2. Lopes P.C., Block P., König B. 2016 Infection-induced behavioural changes reduce connectivity and the potential for disease spread in wild mice contact networks. Sci Rep 6, 31790.

3. Van Kerckhove K., Hens N., Edmunds W.J., Eames K.T. 2013 The impact of illness on social networks: implications for transmission and control of influenza. Am J Epidemiol 178(11), 1655–1662.

4. Farine D. 2017 The dynamics of transmission and the dynamics of networks. J Anim Ecol 86(3), 415–418.

5. Ripperger S.P., Carter G.G., Page R.A., Duda N., Koelpin A., Weigel R., Hartmann M., Nowak T., Thielecke J., Schadhauser M., et al. 2020 Thinking small: next-generation sensor networks close the size gap in vertebrate biologging. PLoS Biol 18(4), e3000655.

6. Stockmaier S., Bolnick D.I., Page R.A., Carter G.G. (in press) Sickness effects on social interactions depend on the type of behaviour and relationship. J Anim Ecol.

7. Stockmaier S., Bolnick D.I., Page R.A., Carter G.G. 2018 An immune challenge reduces social grooming in vampire bats. Anim Behav 140, 141–149.

8. Ripperger S.P., Carter G.G., Duda N., Koelpin A., Cassens B., Kapitza R., Josic D., Berrío-Martínez J., Page R.A., Mayer F. 2019 Vampire bats that cooperate in the lab maintain their social networks in the wild. Curr Biol 29(23), 4139-4144. e4134.

9. Duda N., Nowak T., Hartmann M., Schadhauser M., Cassens B., Wägemann P., Nabeel M., Ripperger S., Herbst S., Meyer-Wegener K., et al. 2018 BATS: Adaptive ultra low power sensor network for animal tracking. Sensors 18(10), 3343.

10. O’Mara M.T., Wikelski M., Dechmann D.K. 2014 50 years of bat tracking: device attachment and future directions. Methods Ecol Evol 5(4), 311–319.

11. Farine D.R. 2017 A guide to null models for animal social network analysis. Methods Ecol Evol 8(10), 1309–1320.

12. Stockmaier S., Bolnick D., Page R., Josic D., Carter G. (in review) An immune challenge reduces contact calling in vampire bats.

13. Carter G.G., Wilkinson G.S. 2016 Common vampire bat contact calls attract past food-sharing partners. Anim Behav 116, 45–51.

14. Warren H.S., Fitting C., Hoff E., Adib-Conquy M., Beasley-Topliffe L., Tesini B., Liang X., Valentine C., Hellman J., Hayden D. 2010 Resilience to bacterial infection: difference between species could be due to proteins in serum. J Infect Dis 201(2), 223–232.

15. Deak T., Bellamy C., Bordner K.A. 2005 Protracted increases in core body temperature and interleukin-1 following acute administration of lipopolysaccharide: implications for the stress response. Physiol Behav 85(3), 296–307.

16. Behringer D.C., Butler M.J., Shields J.D. 2006 Avoidance of disease by social lobsters. Nature 441(7092), 421–421.

17. Kiesecker J.M., Skelly D.K., Beard K.H., Preisser E. 1999 Behavioral reduction of infection risk. P Natl Acad Sci USA 96(16), 9165–9168.

18. Boillat M., Challet L., Rossier D., Kan C., Carleton A., Rodriguez I. 2015 The vomeronasal system mediates sick conspecific avoidance. Curr Biol 25(2), 251–255.

19. Poirotte C., Massol F., Herbert A., Willaume E., Bomo P.M., Kappeler P.M., Charpentier M.J. 2017 Mandrills use olfaction to socially avoid parasitized conspecifics. Sci Adv 3(4), e1601721.

20. Cremer S., Pull C.D., Fürst M.A. 2018 Social immunity: emergence and evolution of colony-level disease protection. Annu Rev Entomol 63, 105–123.

21. Wang C., Liu L., Hao X., Guo H., Wang Q., Huang J., He N., Yu H., Lin X., Pan A. 2020 Evolving Epidemiology and Impact of Non-pharmaceutical Interventions on the Outbreak of Coronavirus Disease 2019 in Wuhan, China. medRxiv. (DOI:https://doi.org/10.1101/2020.03.03.20030593).

22. Ezenwa V.O., Archie E.A., Craft M.E., Hawley D.M., Martin L.B., Moore J., White L. 2016 Host behaviour–parasite feedback: an essential link between animal behaviour and disease ecology. Proc Roy Soc B BiolSci 283(1828), 20153078.

23. Klein S.L. 2003 Parasite manipulation of the proximate mechanisms that mediate social behavior in vertebrates. Physiol Behav 79(3), 441–449.

24. Blackwood J.C., Streicker D.G., Altizer S., Rohani P. 2013 Resolving the roles of immunity, pathogenesis, and immigration for rabies persistence in vampire bats. P Natl Acad Sci USA 110(51), 20837–20842.

25. Montiglio P.-O., Gotanda K.M., Kratochwil C.F., Laskowski K.L., Farine D.R. 2020 Hierarchically embedded interaction networks represent a missing link in the study of behavioral and community ecology. Behav Ecol 31(2), 279–286.

