## Supplementary material for "Sickness behaviour reduces network centrality in wild vampire bats": Figures S1 & S2 (from another version of this preprint: version 1, 2020-03-31)

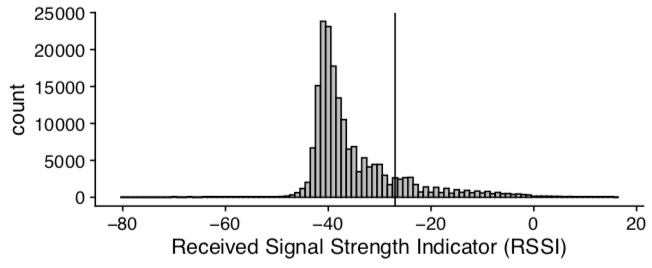

**Figure S1. Threshold of proximity sensor signal strength used to define association.** Vertical line marks the 85% threshold at -27 dBm, which we used for defining association rates in social networks.

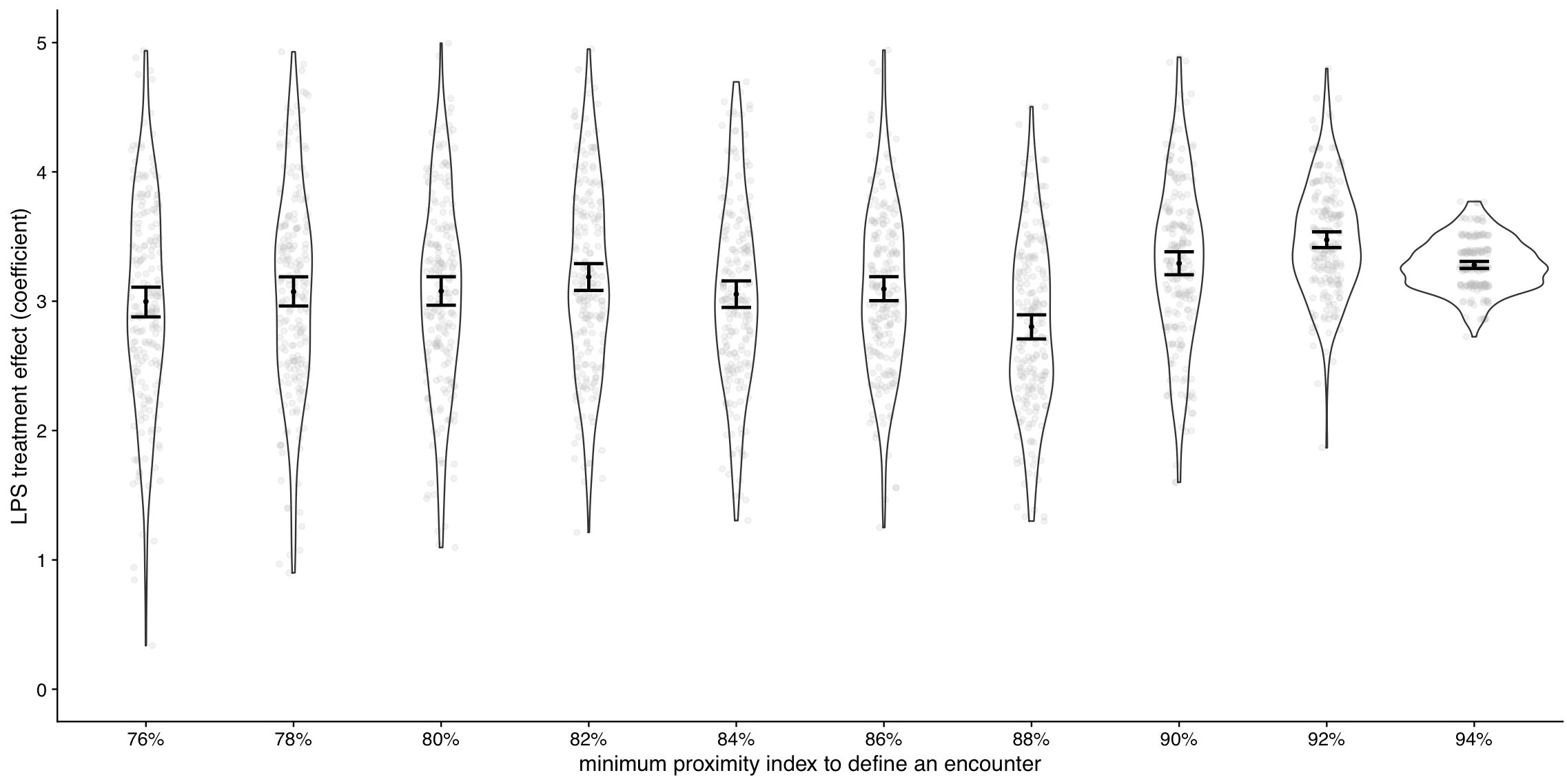

**Figure S2. Stability of the effect size of the LPS treatment on degree centrality across different proximity thresholds and a constant sample size of bats and observations (n=5,258).** The model coefficient for the treatment effect on degree centrality was relatively stable when we randomly sampled 5258 observations that were above different minimum proximity index thresholds (from only the top 24% to only the top 6% of signal intensities).
